# Apomixis-related genes identified from a coexpression network in *Paspalum notatum*, a Neotropical grass

**DOI:** 10.1101/369280

**Authors:** Fernanda A. de Oliveira, Bianca B. Z. Vigna, Carla C. da Silva, Alessandra P. Fávero, Frederico de P. Matta, Ana L. S. Azevedo, Anete P. de Souza

**Affiliations:** Centro de Biologia Molecular e Engenharia Genética (CBMEG), Universidade Estadual de Campinas (UNICAMP), Campinas, SP, Brazil; Embrapa Pecuária Sudeste, São Carlos, SP, Brazil; Embrapa Gado de Leite, Juiz de Fora, MG, Brazil; Departmento de Biologia Vegetal, Instituto de Biologia, UNICAMP, Campinas, SP, Brazil

## Abstract

Apomixis is a highly desirable trait in modern agriculture, due to the maintenance of characteristics of the mother plant in the progeny. However, incorporating it into breeding programs requires a deeper knowledge of its regulatory mechanisms. *Paspalum notatum* is considered a good model for such studies because it exhibits both sexual and apomictic cytotypes, facilitating the performance of comparative approaches. Therefore, we used comparative transcriptomics between contrasting *P. notatum* cytotypes to identify novel candidate genes involved in the regulation of the expression of this phenotype. We assembled and characterized a transcriptome from leaf and inflorescence from apomictic tetraploids and sexual diploids/tetraploids of *P. notatum* accessions, and then assembled a coexpression network based on pairwise correlation between transcripts expression profiles. We identified genes exclusively expressed in each cytotype and differentially expressed genes between pairs of cytotypes. Gene ontology enrichment analyses were performed for the interpretation of data. We *de novo* assembled 114,306 of reference transcripts. 536 novel candidate genes for the control of apomixis were detected through statistical analyses of expression data, contains in this set, the interactions among genes potentially linked to the apomixis-controlling region, differentially expressed, several genes also already reported in the literature and their neighbors transcriptionally related in the coexpression network. The reference transcriptome obtained in this study represents a robust set of expression data for *P. notatum*. Additionally, novel candidate genes identified in this work represent a valuable resource for future grass breeding programs.

**Author Summary:** Clonal mode of reproduction by seeds is termed apomixis, which results from the failure of gamete formation (meiosis) and fertilization in the sexual female reproductive pathway. The manipulation of seeds production genetically identical to the mother plant bears great promise for agricultural applications, however clarification regarding gene interactions involved in reproductive process is needed. *Paspalum* is considered a model genus for the analysis of apomixis mechanisms. Here, we describe an overall analysis of the expression profiles of *Paspalum notatum* transcripts in response to changes in reproductive mode (sexual to apomictic), which allowed us to identify several candidate apomixis genes. Among these, we found genes potentially associated with the apomixis control region, in addition to genes already described in the literature for *Paspalum*, which highlights the representativeness of assembled transcriptome. For the first time in the literature, we explored the main biological processes involved in controlling the expression of apomictic reproduction based on co-regulatory networks of candidate apomixis genes.

## Introduction

Apomixis is a mode of asexual reproduction through seeds in which the plants produced offspring is genetically identical to the female parent [1]. In apomixis, a non-reduced cell undergoes developmental pathways different from those of sexual cells. The multiple trajectories are classified as sporophytic or gametophytic according to the developmental origin of the cell from which the embryo is derived [2-4]. In sporophytes, the embryo develops from a somatic cell of the ovule through numerous mitotic divisions. In gametophytic apomixis, a chromosomally unreduced embryo sac can be formed either from the megaspore mother cell (diplospory) or from a nearby nucellar cell (apospory) without fertilization (parthenogenesis) through a process termed apomeiosis. Endosperm formation is required to produce a viable seed [5,6]. Apomixis is widely distributed among angiosperms, among which Poaceae represents the family with the largest number of apomictic genera, with the 47 apomictic species of *Paspalum* standing out in particular [7-9].

The polyploid nature of all apomicts provides a challenge for genetic and genomic analysis [10]. Polyploidization is known to cause immediate and extensive genomic changes, including sequence rearrangements and/or elimination, changes in DNA methylation, and the loss of balance in gene expression [11,12]. Apomixis is frequently correlated with polyploidy and might have arisen through deregulation of the sexual developmental pathway, due to the increase in number of genomic complements, through a mechanism regulated by both genetic and epigenetic components [13-16].

*Paspalum notatum* Flüggé, also known as bahiagrass, belongs to the genus *Paspalum,* which exhibits numerous characteristics that make it a very interesting system for the study of apomixis [8]. This species is considered an agamic complex that includes different ploidy levels and reproductive modes in which diploid cytotypes (2n=2×=20) are sexual and self-incompatible, whereas polyploids (3×, 4×, 5×, and 6×) are self-compatible pseudogamous apomicts [17]. No 4× sexual cytotypes have been found in nature, but they have been obtained artificially [18,19]. The inheritance of apomixis in *Paspalum* is controlled by a single complex dominant locus that confers apospory, i.e., epigenetically controlled parthenogenesis [20], with the capacity to form endosperm with a maternal excess genome contribution ratio of 4:1 (maternal:paternal) [8]. The apomixis-controlling region (ACR) is small compared to other apomictic systems [8], does not demonstrate recombination and conserves a relatively narrow region that is linked to apomixis among the distinct species [21,22]. Comparative mapping of the ACR has shown synteny with a portion of the long arm of chromosome 12 of rice in *P. simplex, P. malacophyllum* and *P. procurrens* [21-23]. In *P. notatum* the ACR shows synteny with regions of rice chromosomes 2 and 12, suggesting the presence of chromosomal rearrangements [21, 24-26].

Apomixis presents potential significance for agriculture, allowing the maintenance of heterosis in hybrid progeny. Therefore, this mode of reproduction has become the subject of exhaustive cytoembryological, cytogenetic and molecular analyses [10,25]. Some key genes associated with the components of apomixis and the isolation of some sequence candidates have already been discovered [6,12,14,27-41]. However, insights into the genetic mechanisms underlying asexual reproduction in natural apomicts are still needed to develop a stable and universal apomixis system to be applied in breeding programs [39]. Given the complexity of this trait, an understanding of the genomic structure of the apomictic locus is likely to be an essential prerequisite for manipulation of the sexual pathway in model plants or economically important crops [15]. However, the suppression of recombination events around the ACR limits forward genetic strategies to isolate the mechanism triggering apomixis by map-based [31]. In this context, identify and validate genes that are differentially expressed in apomicts and to conduct more meticulous investigations aiming to detect their effects in phenotype, have garnered extreme interest in the study of apomixis [31,40].

RNA sequencing (RNA-seq) is the most effective method for simultaneously predicting new transcripts and identifying differentially expressed genes in distinct tissues, genotypes, abiotic conditions or developmental stages [42,43]. Conversely, considering the large amount of data generated from RNA-seq, new approaches that efficiently extract meaningful associations from highly multivariate datasets are needed [44]. The construction of coexpression networks from gene expression data using pairwise correlation metrics provides us with valuable information on alterations in biological systems in response to differential gene expression patterns [44-46].

The objective of this study was to identify candidate genes possibly involved in the regulation of the expression of apomixis in *P. notatum*. To reach this goal, we used RNA-seq technology to obtain a reference transcriptome and comprehensively analyze gene expression profiles in leaves and florets from apomictic and sexual genotypes with different ploidy levels in *P. notatum*. Our study revealed several differentially expressed genes among the analyzed genotypes, and a coexpression network analysis and Gene Ontology (GO) annotation of transcripts allowed us to investigate the main biological processes of candidate genes potentially linked to the ACR. These candidate genes may be used to further explore and clarify the mechanisms regulating apomixis in forage grasses.

## Results

### Verification of the DNA content and mode of reproduction of *P. notatum* accessions

The cytoembryological analysis confirmed the expected mode of reproduction of the accessions based on the literature (Tables 1 and 2). The 2C DNA content of all plants from accessions BGP_22 and BGP_306 was half the 2C DNA content from all plants from BGP_30, BGP_34, BGP_115, and BGP_216. Therefore, all evaluated plants had DNA contents compatible with the described ploidy levels of their respective accessions in the literature (Tables 1 and 2).

**Table 1.**
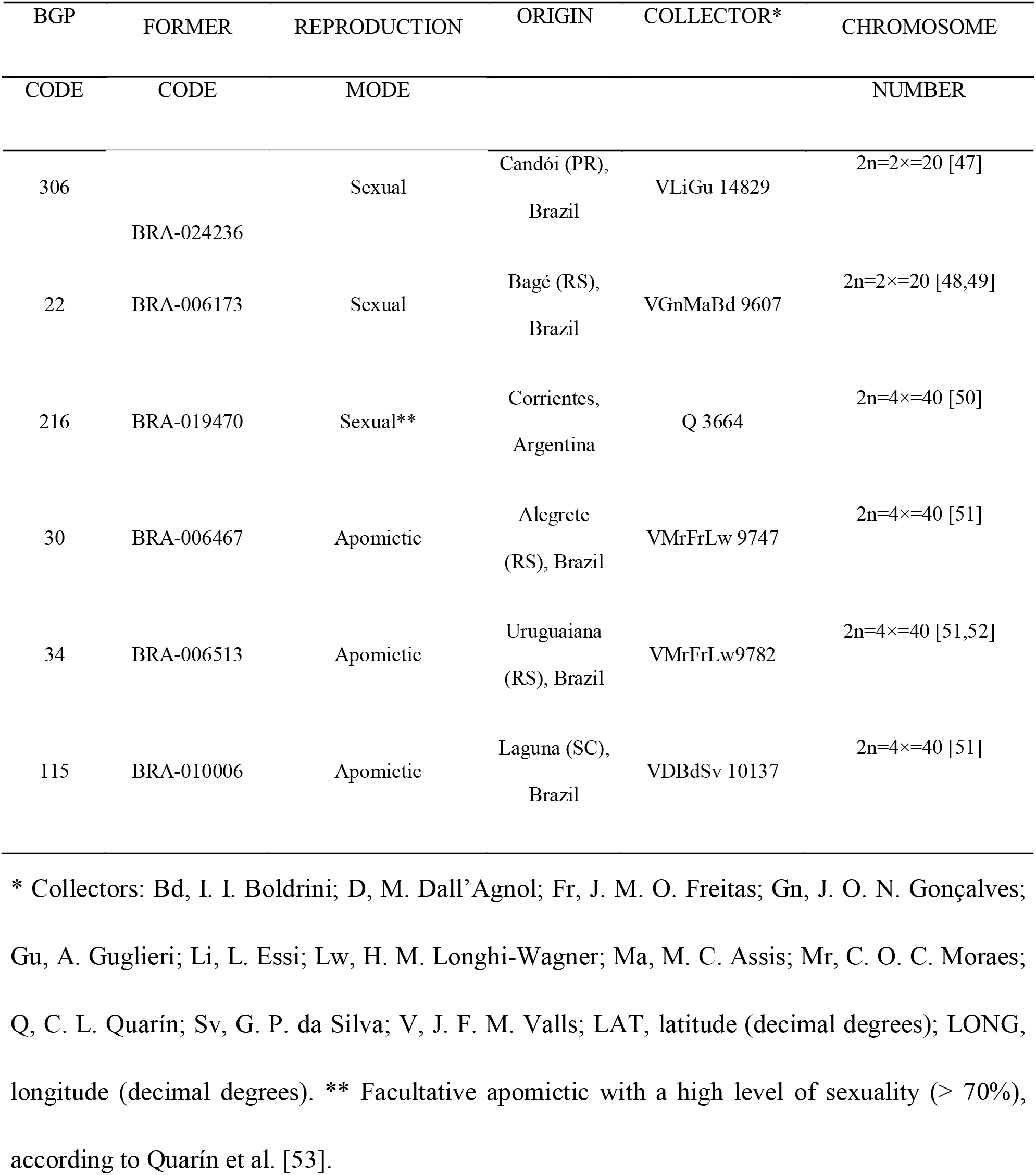
*Paspalum notatum* Accessions Evaluated in this Study.

**Table 2.**
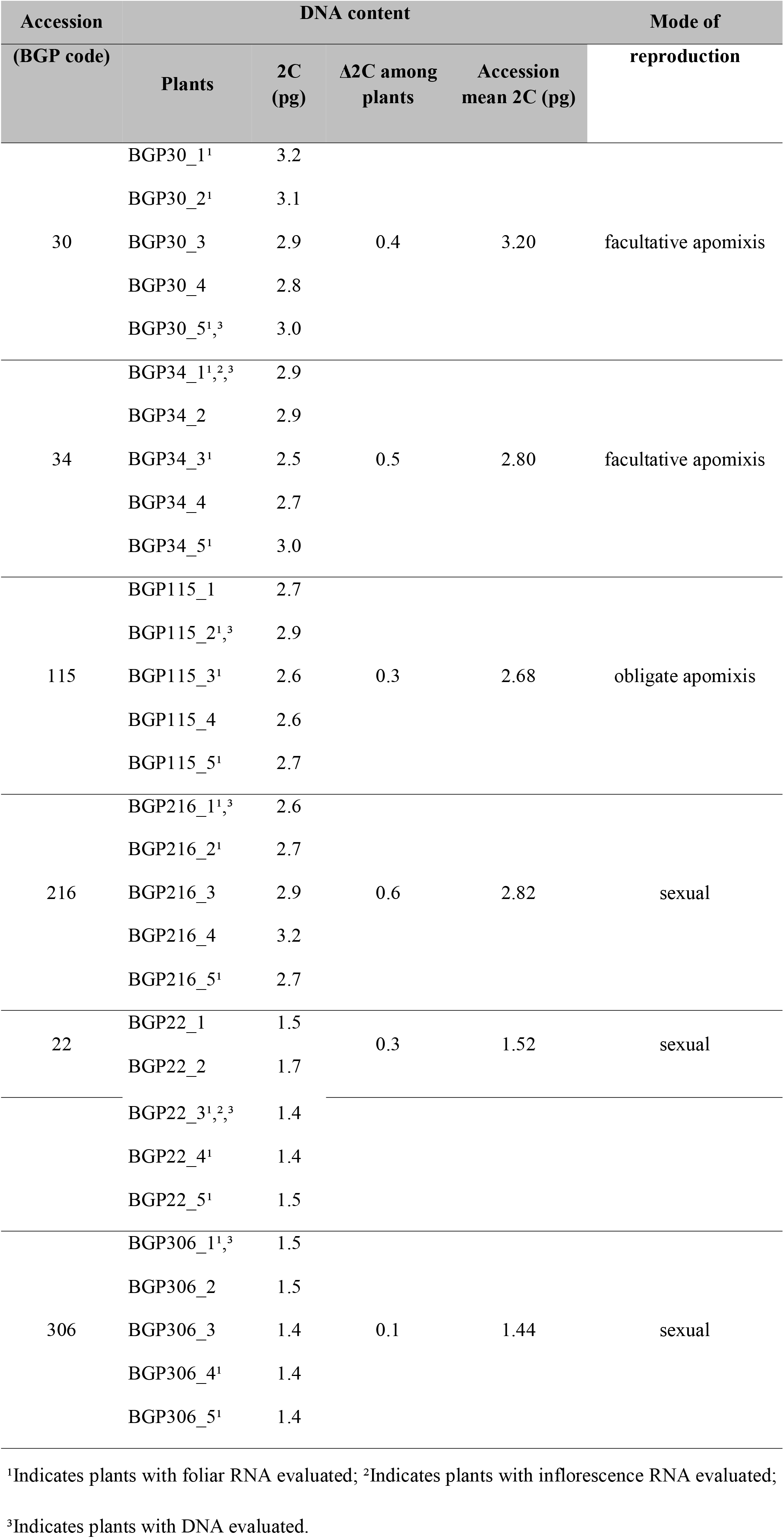
DNA Contents and Modes of Reproduction of the Bahiagrass Accessions Evaluated in this Study.

### RNA-seq analysis and *de novo* transcriptome assembly

After removing barcodes sequences and low-quality and contaminant reads, 313,198,097 high-quality 72-bp paired-ends reads from floret and leaf tissues were used to assemble a *P. notatum* reference transcriptome. A total of 203,808 transcripts were assembled, out of which a set of 114,306 non-redundant transcripts were filtered (56.08% of all transcripts) (Table 3). Final transcripts in the reference transcriptome had an average length of 750.51 bp with an N50 of 906 bp and GC percentage of 47.60%. According to the length distribution of non-redundant transcripts, we obtained 21,585 (18.88%) transcripts that were longer than 1 kb, a size range that commonly confers a high annotation rate. More than half of the total annotated transcripts were > 500 bp in length (Fig S1).

The Bowtie aligner mapped 94.39% of sequenced reads onto the assembled transcripts. 73.78% mapped to the non-redundant transcripts set. Out of these, 82.97% and 83.48% of the sequencing reads belonged to sexual and apomictic samples, respectively, showing similar representation within the final transcriptome. The BUSCO analysis included 956 conserved single-copy plant orthologs; the *P. notatum* transcriptome showed high assembly completeness, with 664 (69%) complete sequences (532 as a single copy and 132 as multiple copies) and 169 (17%) fragmented sequences. One hundred and twenty-three (12%) BUSCO plant orthologs were not identified in the reference *P. notatum* transcriptome.

**Table 3.**
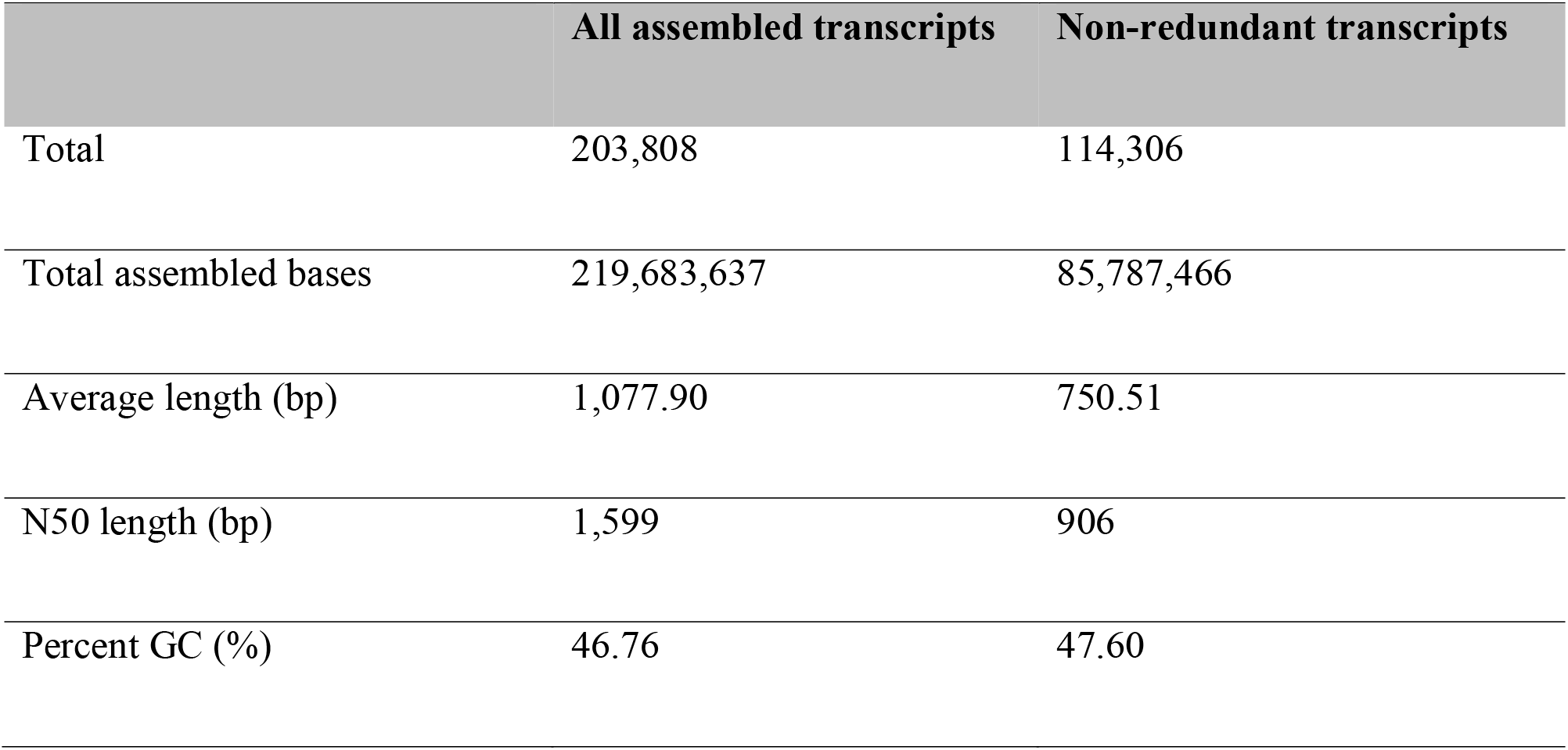
Statistics for the Assembled Transcriptome of *Paspalum notatum*.

|  | All assembled transcripts | Non-redundant transcripts |
| --- | --- | --- |
| Total | 203,808 | 114,306 |
| Total assembled bases | 219,683,637 | 85,787,466 |
| Average length (bp) | 1,077.90 | 750.51 |
| N50 length (bp) | 1,599 | 906 |
| Percent GC (%) | 46.76 | 47.60 |

### Transcriptome annotation

Among the 114,306 unigenes, 51,939 (45.44%) were similar to proteins from the NCBI Nr database, 34,750 (30.40%) to proteins from the UniProtKB/Swiss-Prot, and 54,568 (47.74%) to proteins from Phytozome.

The GO database was then used to retrieve the GO identifiers for all possible transcripts. Overall, 21,537 (41.47%) transcripts were assigned to 4,614 GO terms in three main categories: 2,510 biological process, 1,486 molecular function, and 618 cellular components. The KEGG annotation was possible for 9,221 annotated transcripts belonging to 129 pathways. Among these, the most represented were Purine Metabolism Pathway (1,079 members), followed by Metabolism of Thiamine (907 members) and Starch and Sucrose Metabolism (336 members). Overall, 33,873 transcripts remained without hits after searching against all protein databases. We then performed a search for the presence of open reading frames (ORFs), which were found in 15,107 transcripts, 2,347 of which were complete. These sequences that showed ORFs without annotation require further investigation since they may represent genes that have not yet been described and possibly new genes that may be unique to *P. notatum*.

### Expression levels of genes and identification of DEGs

The sequenced reads were realigned against the 114,306 transcripts of the reference transcriptome to estimate the expression level of the transcripts in each genotype. Most identified transcripts had expression levels of FPKM ≤ 100 (110,621 transcripts), whereas only 49 transcripts had high expression levels (FPKM ≥ 400), in these, most of them involved in photosynthesis process, house-keeping genes and response to stress. The remaining transcripts were not mapped and thus, their expression levels could not be determined. A comparison of the 110,656 transcripts that showed different levels of expression in the two tissues of the sequenced samples revealed genes that were expressed in the genotypes of only one phenotypic class and that were not expressed in the others (Fig 1), which were considered "unique" transcripts. Thus, 19,304 unique transcripts were found exclusively in the 4×APO group, 2,173 in the 2×SEX group, and 217 in the 4×SEX group. All transcripts expressed in florets were also expressed in leaves; there were no unique transcripts expressed in the florets from either diploid sexual or tetraploid apomictic samples (Fig 1).

**Fig 1.**
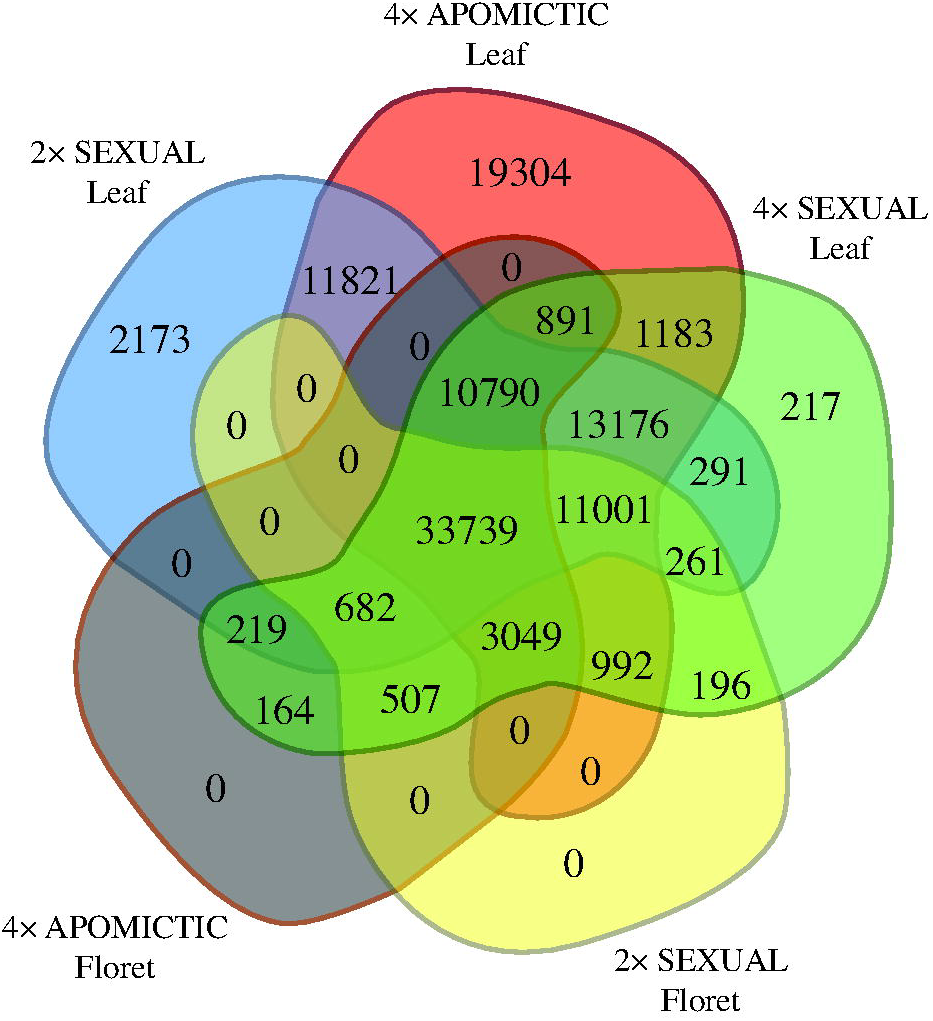
Venn diagram showing the distribution of *P. notatum* transcripts. Expression level of transcripts from two tissues (floret and leaf) and each phenotypic class (sexual diploid, sexual tetraploid, and apomictic tetraploid).

For pairwise differential expression analyses of all three phenotypic classes, we considered only leaf samples because they expressed all representative transcripts from the transcriptome assembly and had three clones for each genotype, maximizing the accuracy and reliability of the differential gene expression analysis. However, within the same phenotypic class, different expression patterns were observed among genotypes, mainly in 4×APO samples (Fig S2). Thus, to reduce the effects of the differences in expression caused by genotype, we performed a two-step pipeline. In the first step, we identified 72,318 transcripts that were equally expressed among all 4×APO genotypes and 47,069 equally expressed transcripts in the 2×SEX genotypes. This initial analysis was not necessary for the 4×SEX because only one genotype was analyzed. From these, we selected 28,969 transcripts that were expressed in all three phenotypic classes. In the second step of the differential expression analysis, pairwise comparisons of 28,969 transcripts allowed the identification of 2,072 DEGs between 2×SEX and 4×APO, which included 772 and 1,173 overexpressed transcripts, respectively. There were 1,173 DEGs between 2×SEX and 4×SEX, including 308 and 798 overexpressed transcripts, respectively, and 1,317 were identified as DEGs between 4×APO and 4×SEX, 618 and 661 of which were overexpressed, respectively. Next, we inferred the enriched functions for the identified DEGs that were over- or underexpressed (Figs S5-S7).

Among the identified DEGs, we isolated a set of 510 transcripts that were consistently differentially expressed between apomictic and sexual samples independently of the ploidy level (diploid or tetraploid). We identified GO terms enriched in this set of transcripts, which belonged to 26 biological processes (BP), 13 molecular functions (MF), and three cellular components (CC) (Fig 2). We considered these transcripts as potentially involved in the determination of the reproductive mode.

**Fig 2.**
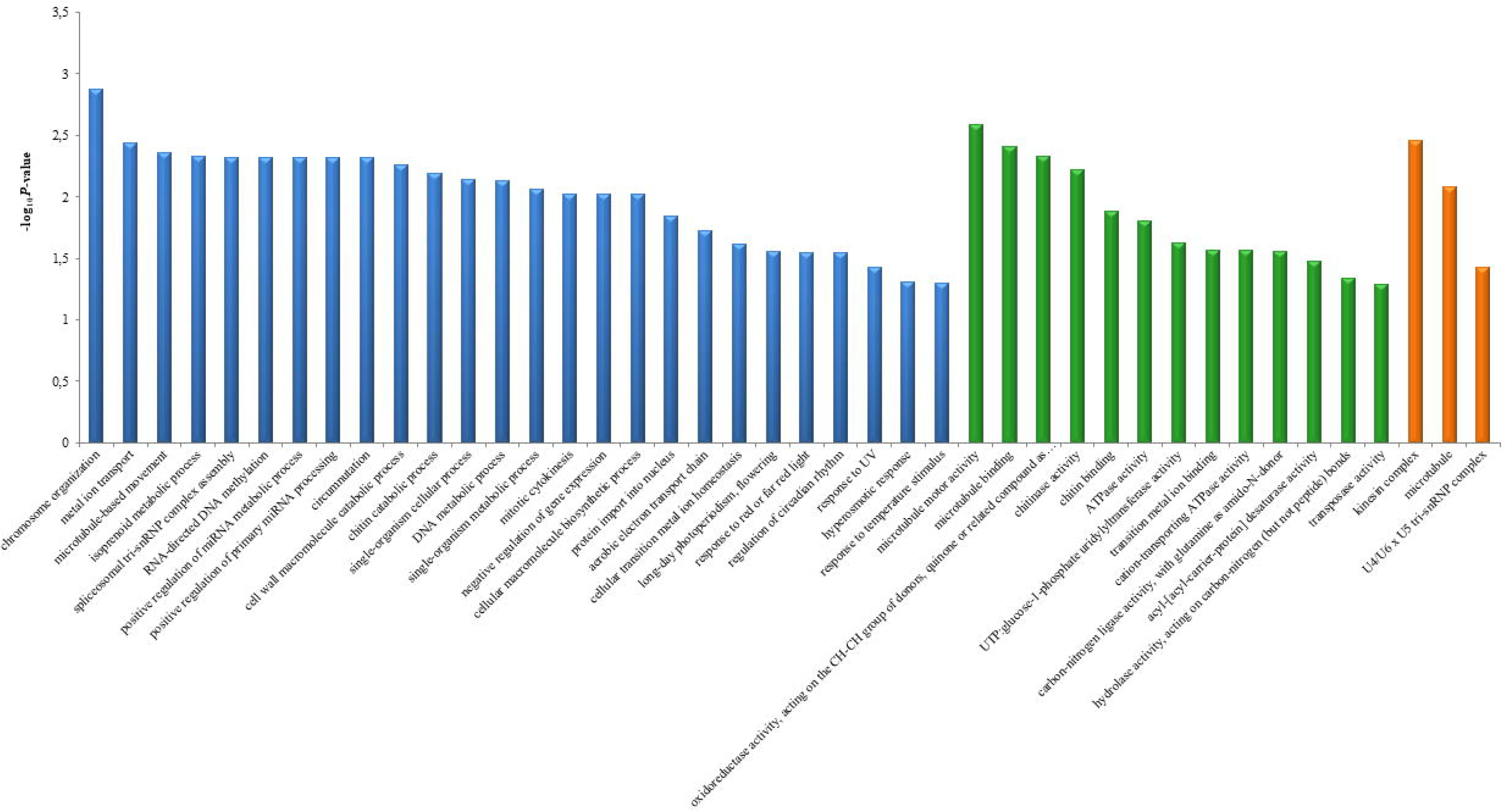
Enriched ontology terms of 510 transcripts differentially expressed in the *P. notatum* transcriptome. Transcripts consistently differentially expressed between apomictic and sexual samples.

### qRT-PCR validation of RNA-seq data

We selected a subset of 18 DEGs detected in-silico between apomictic and sexual samples to confirm the expression profiles through an independent technique, qRT-PCR. Five equally expressed transcripts detected in-silico showed desirable expression stability and coefficient of variance in qRT-PCR reactions (Table S1), out of which one was selected to be an internal control for qRT-PCR verification of 2×SEX vs. 4×APO analyses (01RefGen-*Pnot*) and one for the verification of 4×APO vs. 4×SEX analyses (04RefGen-*Pnot*).

Out of 18 primer pairs designed for target DEGs, eight (Table S2) were used for qRT-PCR validation of RNA-seq data. The qRT-PCR analyses revealed consistent results compared to those detected through RNA-seq (Figs S3 and S4), emphasizing the power of the latter technique. Two primers pairs were not evaluated in qRT-PCR but showed interesting results in terms of DNA amplification. The first was designed from a transcript that was exclusively expressed in apomict samples in RNA-seq analyses (i.e., showing a zero expression value in the sexual samples). This transcript did not amplify using genomic DNA from sexual samples as PCR templates. For the second primer pair, although genomic DNA amplification occurred in all samples, when complementary DNA (cDNA) was used as PCR template, only apomictic samples presented amplicons. The expression level of this transcript estimated from RNA-seq of the same samples was very low in 2×SEX samples. These results suggest that the first transcript, expressed only in apomictic cytotypes, is exclusive to the genomes of these samples, whereas the second transcript was poorly expressed or silenced in sexual samples.

### Detailed search for potential apomixis genes

We detected 1,612 transcripts that showed high similarity to the ACR in the *P. notatum* reference transcriptome. Out of these, 1,356 transcripts aligned to corresponded rice chromosome 2, and 256 aligned to rice chromosome 12. Curiously, 40 of these transcripts were part of the set identified as unique to apomictic samples, nine were unique to sexual diploids, and six were unique to sexual tetraploid samples expression. Moreover, considering the analyses of differential expression, 16 putative ACR transcripts were differentially expressed in 2×SEX vs. 4×APO, 20 DEGs in 4×APO vs. 4×SEX, and 17 DEGs in 2×SEX vs. 4×SEX. All these findings, especially this set of 108 unique or differentially expressed transcripts, represent a valuable set of genes that deserve to be carefully investigated to determine their roles in apomixis and determine whether they are effectively genetically within the ACR in *P. notatum*.

The BLAST results in Table S3 show the similarity scores of the *P. notatum* transcripts to previously *Paspalum* sequences associated to the apomictic mode of reproduction, revealing that our transcriptome contains genes already found to be involved in asexual reproductive development in *Paspalum*.

### Transcripts coexpression network

We identified 879,481 connections among 53,262 transcripts from RNA-seq data (Fig 3). Transcripts were grouped into 642 clusters according to their correlated pattern of expression level. In the network, the unique transcripts from each phenotypic class tended to form coexpression modules (Fig 3B-D). Despite this subdivision, the relevant biological relationships of these transcripts with all remaining transcripts can be recovered in a fully integrated network.

**Fig 3.**
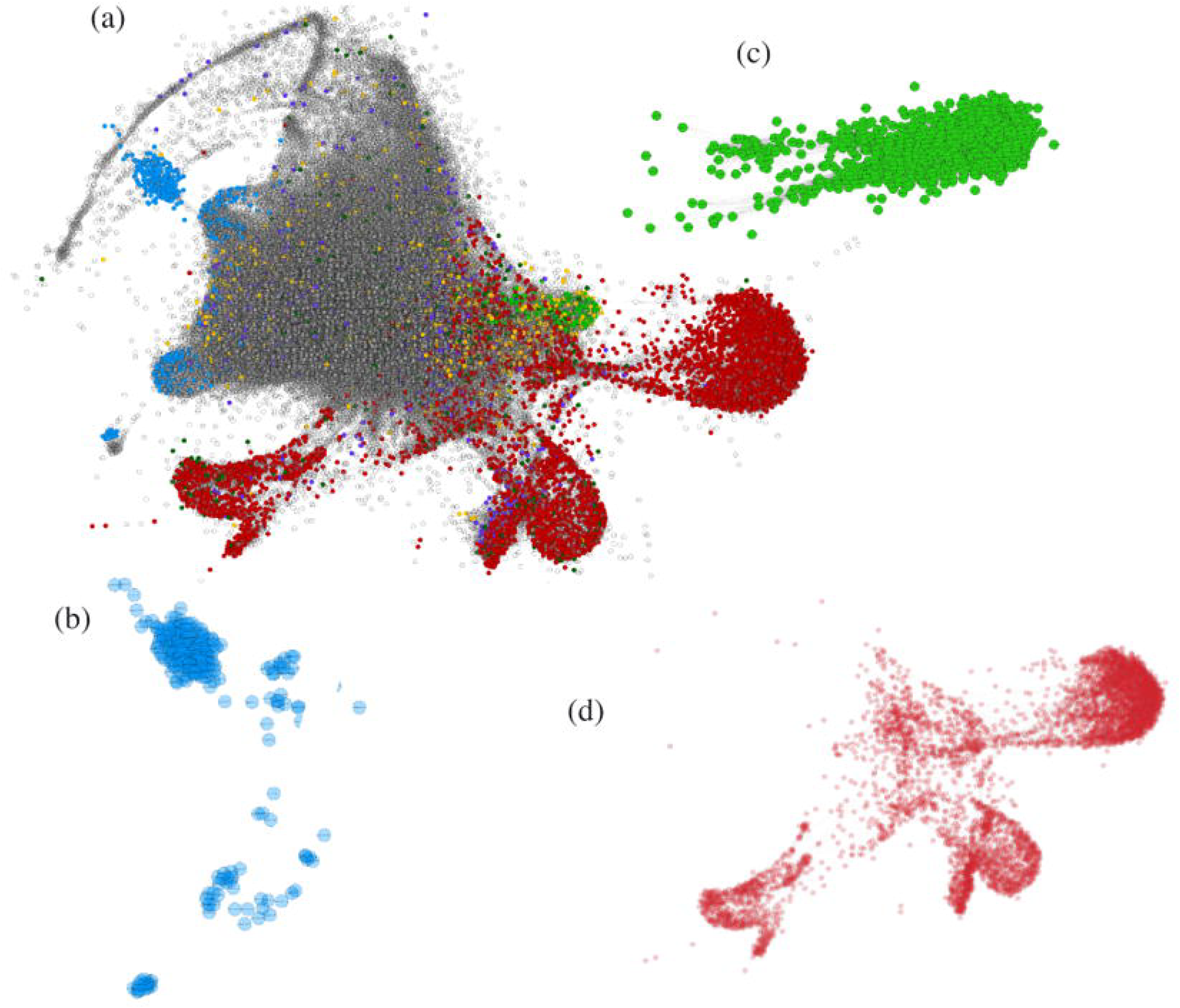
Coexpression network for *Paspalum notatum*. (A) Coexpression network of all analyzed transcripts; the more centralized genes in light gray are those common to all genotypes analyzed with non-differential expression. Node color denotes the differentially expressed transcripts between the phenotypic classes: purple (2×SEX vs. 4×APO), yellow (2×SEX vs. 4×SEX), and dark green (4×APO vs. 4×SEX). The networks of exclusively expressed transcripts for each phenotypic class are highlighted: (B) transcripts unique to sexual diploids (blue); (C) transcripts unique to sexual tetraploids (green); and (D) transcripts unique to apomictic tetraploids (red). Edges denote interaction strength. Circular nodes represent transcripts.

We identified a direct correlation among the 108 differentially expressed and/or unique putative ACR genes and their first neighbors in the gene expression network. Thus, we retrieved a sub-network composed of 536 strongly correlated transcripts (Fig 4). We used GO enrichment analysis to summarize the main putative functions of this set of transcripts, which included 43 biological processes (BP), 31 molecular functions (MF) and 17 cellular components (CC) (Fig. 5).

**Fig 4.**
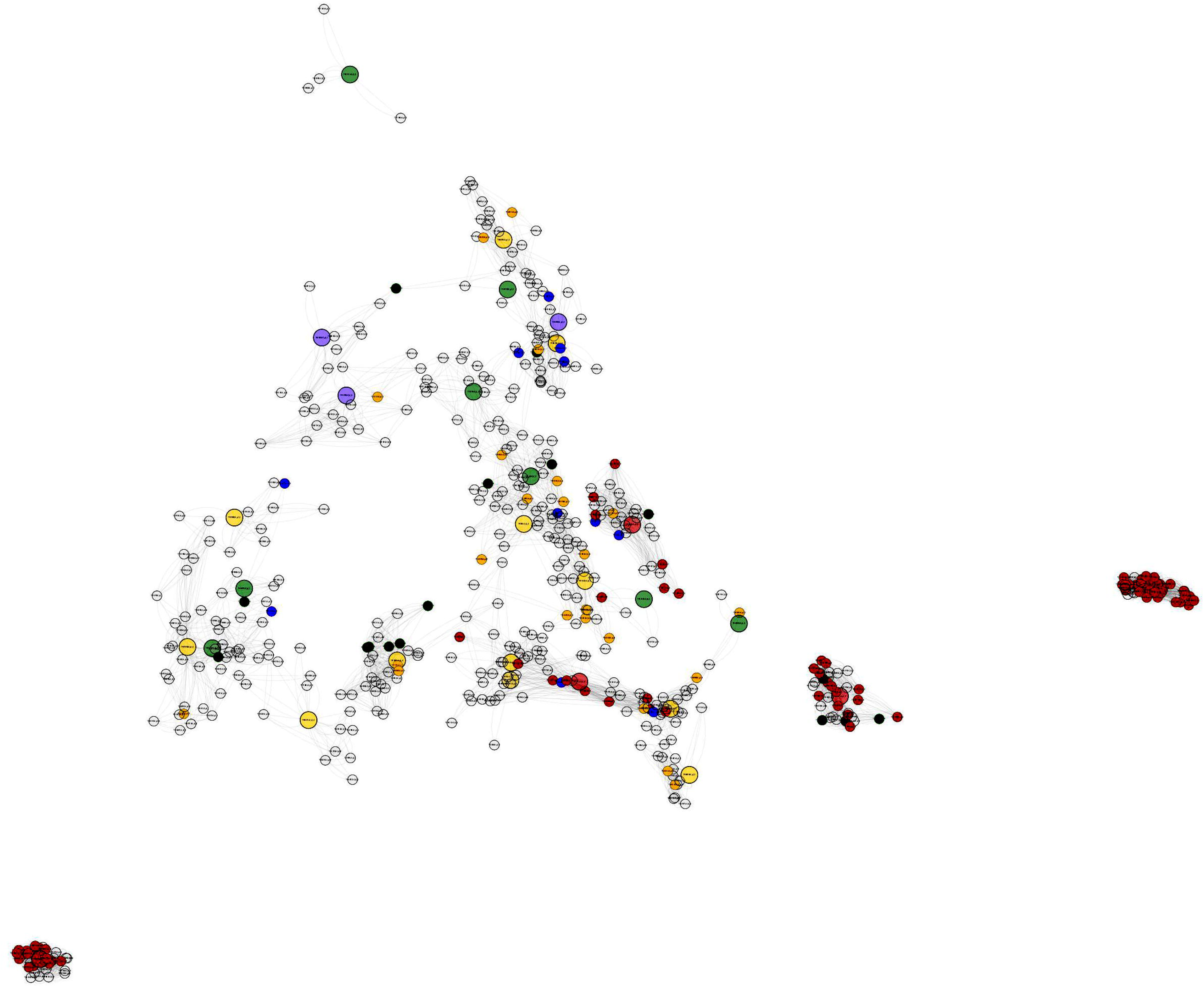
Gene expression sub-network for 536 *Paspalum notatum* transcripts. The 108 differentially expressed and/or unique putative ACR transcripts are presented in this sub-network, along with their first neighbors. The color patterns are the same as those used in the complete coexpression network in Fig 3.

**Fig 5.**
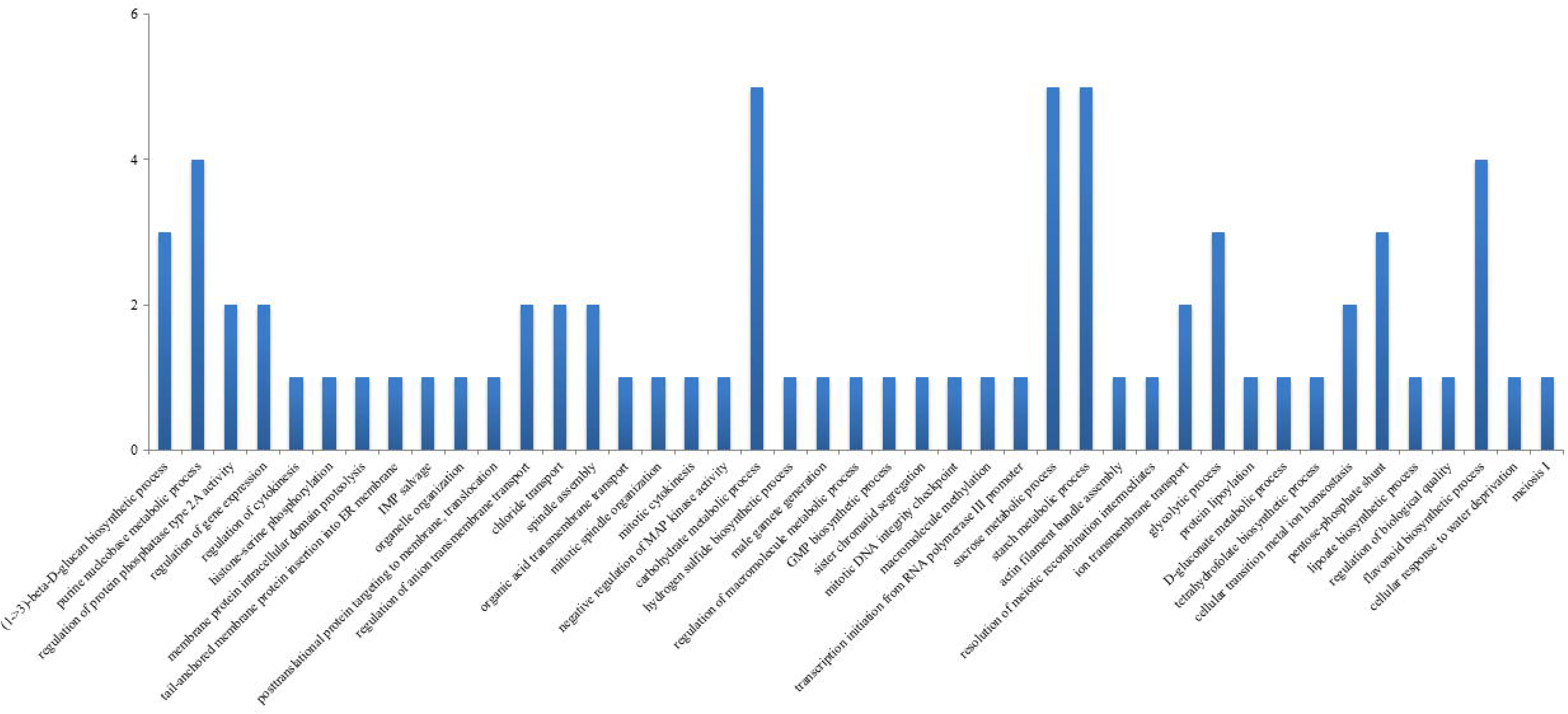
Enriched biological processes in the 536 transcripts in the gene expression sub-network.

## Discussion

We constructed a *P. notatum* reference transcriptome using the next-generation sequencing approaches, RNA-seq, to investigate the gene expression changes associated with apomictic and sexual reproduction. RNA-seq enabled the assembly of a non-redundant transcriptome, containing 114,306 transcripts from florets and leaves from six distinct genotypes with different ploidy levels and reproductive modes. Robust metrics indicated that the transcriptome was a quality assembly with a high degree of integrity, which further expands the bahiagrass transcriptome database.

Even though we did not sequence inflorescences samples at different developmental stages (premeiosis, late premeiosis/meiosis, postmeiosis and anthesis) to search for candidate apomixis-linked genes, differential expression analyses between leaf samples from sexual and apomictic cytotypes allowed us to identify DEGs that may be representative of the mode of reproduction and not dependent of the level of ploidy. We isolated expression patterns of phenotypic classes, aimed both at removing transcripts whose expression could be related to the effect of genotype and at identifying transcripts expressed in all genotypes within the same phenotypic class.

GO classification of the assembled transcripts into 4,614 known terms, was used to perform functional enrichment analyses of candidate apomixis genes. Based on a set of 510 DEGs detected between apomictic and sexual, among the enriched GO terms, we emphasize “positive regulation of miRNA metabolic process (GO:2000630)” and “regulation of primary miRNA processing (GO:2000636)”, both of which either activate or increase the frequency, rate, or extent of miRNA production. miRNA, in turn, is directly related to “DNA metabolic process (GO:0006259)”, which decreases the rate of gene expression (negative regulation of gene expression - GO:0010629) through an epigenetic RNA-based gene silencing process (RNA-directed DNA methylation - GO:0080188). Five transcripts were mainly involved in these processes and showed higher expression levels in apomict samples in comparison to sexual samples; these transcripts correspond to a pseudoARR-B transcription factor; pre-mRNA splicing factor, putative (*Ricinus communis*); hypothetical protein SORBIDRAFT_05g016770; uncharacterized protein, LOC100501330 (*Zea mays*); and an uncharacterized protein LOC103641690. The GO analysis reveals the significant enrichment of terms associated with the regulation of gene expression by epigenetic silencing among transcripts showing higher expression levels among apomictic samples in comparison to sexual samples of the bahiagrass. This result is consistent with the growing body of evidence that suggests that apomixis arises from deregulation of the sexual pathway, where epigenetic mechanisms play a significant role in at least some elements of apomictic development [8,15,54].

These DEGs represent a set of candidate genes that, together with the 19,304 transcripts exclusively expressed in apomictic samples, deserve further investigation. In apospory, gene expression occurs at specific stages of apomictic development such as apomeiosis, parthenogenesis, and endosperm development. Increased or decreased expression of some genes during these specific stages may hinder the analysis of differential expression between sexual and apomictic genotypes [40,55]. Nonetheless, we believe that the identification of DEGs as performed herein, using selected transcripts with detectable expression pattern among all genotypes of all phenotypic classes, may have minimized this influence. At the same time, our approach increased the potential for discovery of candidate genes involved in apomictic process and not only in a single step. Thus, future reverse genetics experiments based on qRT-PCR and in situ hybridization could be useful to identify the role of specific genes in the whole apomixis process.

Functionally related genes tend to be transcriptionally coordinated [56,57]. Therefore, the construction of transcripts coexpression network provided a powerful resource, for the identification of transcripts that are coexpressed with unique and differentially expressed genes, despite showing undetectable differences in expression between samples. The sub-network containing 536 candidate coexpressed transcripts associated with the expression of apomixis is an example of how we can recover the relevant biological relationships of genes of interest using the transcriptome sequencing approaches. By starting on a smaller scale of transcripts that are DEGs and/or unique genes and possibly integrated into the ACR and combining the information by adding their nearest neighbors, we can obtain a broader view of the processes involved in the regulation of candidate genes. Significantly enriched BP were mainly associated with plant reproduction, for instance, “male gamete generation (GO: 0048232)”; “sister chromatid segregation (GO: 0000819)”; “meiosis I (GO: 0007127)”; “resolution of meiotic recombination intermediates (GO: 0000712)”; “regulation of cytokinesis (GO: 0032465)”; “mitotic cytokinesis (GO: 0000281)”; “spindle assembly (GO: 0051225)”; “mitotic spindle organization (GO: 0007052)”; and “mitotic DNA integrity checkpoint (GO: 0044774)”. In addition, the sub-network was enriched in epigenetic processes such as “macromolecule methylation (GO: 0043414)”; “histone-serine phosphorylation (GO: 0035404)”; “regulation of protein phosphatase type 2A activity (GO: 0034047)”; and “negative regulation of MAP kinase activity (GO: 0043407)”. Transcripts from this sub-network were associated with reproductive processes and the regulation of epigenetic changes by modulating histones. In particular, MAP kinase activity was negatively regulated. The mitogen-activated protein 3-kinase (MAP3K) gene family has been identified as differentially expressed in apomictic and sexual flowers of *P. notatum* [31] and in flowers of *P. simplex* [14], and it might play a role in parthenogenetic development of the embryo in both species [8]. Recently, an essential gene to the formation of unreduced embryo sacs in *P. notatum* was identified from the characterization of a MAP3K retrieved in previous transcriptomic surveys [58].

The size of the ACR in *Paspalum* is relatively small compared with other apomictic systems [8], but the lack of recombination makes it difficult to study this region by map-based cloning [31]. Thus, the discovery of DEGs that are potentially located within this region is extremely valuable for understanding the complex regulatory network of gene–gene interaction. These DEGs could also be used for future manipulation of the apomixis trait, which has outstanding importance in agricultural biotechnology.

In the genus *Paspalum*, the first approach for understanding the molecular basis of apomixis was applied by Pessino et al. [27]. This approach led to the identification of three small sequences that are highly expressed during early megagametophyte development in apomictic plants through differential display experiments in the inflorescences of sexual and aposporous *P. notatum*. Here, we recovered the reported sequences as a single representative transcript (Table S3) similar to a kinesin motor protein, involved in the biological process of microtubule-based movement. The kinesin motor protein has also been reported by other authors as differentially expressed between apomictic and sexual plants [29,31,59]. Interestingly, the transcript assembled in this work also showed differential expression levels between apomictic and sexual samples. Since then, several other candidate genes have been reported using *P. notatum* and *P. simplex* species [14,27,30,31] and more recently, the first approach using RNA-seq was published for bahiagrass based on expression profiling of apomictic and sexual flowers [40]. Some of these candidate genes had been investigated in more detail [35, 37-39]. However, many questions remain unanswered, and further research is needed to define the relationships between the structure, the position, and the function of the known apomixis-linked genes [8]. Furthermore, based on the set of genes selected through differential expression patterns, we recovered previously published sequences in our *P. notatum* transcriptome.

One of the candidate genes previously reported in the literature is n20gap-1, a lorelei-like *P. notatum* gene [31,35] encoding a GPI-anchored protein that supposedly plays a role in the final stages of the apomixis developmental cascade. Laspina et al. [31] previously reported this sequence as linked to the chromosomal locus governing apospory at a genetic distance of 22 cM. Here, one transcript similar to this gene present in the set of transcripts potentially integrated within the ACR, corresponding to rice chromosome 2. Additionally, we found transcripts, including some that were differentially expressed, that aligned with *P. notatum* sequences characterized by Podio et al. [38], corresponding to *Somatic Embryogenesis Receptor-Like Kinase* (*SERK*), in addition to other numerous transcripts with the same annotation. The candidate SERK-like genes play crucial roles in somatic embryogenesis in angiosperms and have been reported as related to apomixis. Albertini et al. [59] analyzed two members of this protein family, namely *PpSERK1* and *PpSERK2*, and found that *PpSERK1* expression levels was high during premeiosis and decreased during the meiosis and post-meiotic stages, whereas *PpSERK2* expression was high from premeiosis to anthesis in the nucellar cells of aposporous genotypes. The authors proposed that the expression pattern of *PpSERK* in *Poa pratensis* was compatible with its role in the specification of aposporous initials. In *P*. *notatum*, the expression of two different members of the *SERK* family (*PnSERK1* and *PnSERK2*) was observed, and *PnSERK2* displayed a spatial expression pattern similar to that reported for the *PpSERKs*, which are expressed in nucellar cells at meiosis in the apomictic genotype [38]. We also identified transcripts that showed similarity to other interesting candidate genes related to apospory, such as the PnTgs1-like gene that encodes a trimethylguanosine synthase-like protein, which plays a fundamental role in nucellar cell fate, as its diminished expression is correlated with initiation of the apomictic pathway in plants [37]. Additionally, two transcripts were similar to the sequence of *Ps*ORC3, which seems to play an active role in mechanisms repressing sexuality in apomictic *P. simplex*, acting in the development of apomictic seeds, which deviate from the canonical 2 (maternal):1 (paternal) genome ratio [39]. The recovery of these previously published sequences demonstrates the representative and informative nature of assembled transcriptome. Indeed, this outcome reaffirms that high-throughput sequencing technology enhances our understanding of global RNA expression.

In this study, we applied RNA-seq technology and bioinformatics methods to assemble a useful transcriptome and identify differences between apomictic and sexual reproduction. Our results reveal DEGs, genes exclusively expressed in apomicts or sexuals, of which are in potential association with the ACR genomic region, including the apo locus. The validation of genes from this set of candidates may enable valuable insights to the understanding of apospory. Moreover, these genes may be considered to screen for molecular markers linked to apomixis in *P. notatum*, which will be crucial to boost breeding programs for apomictic forage grasses.

## Materials and methods

### Plant materials collections and RNA extraction

The six accessions of *P. notatum* used in this study belong to the Germplasm Bank of *Paspalum*, maintained by Embrapa Pecuária Sudeste, São Carlos, SP, Brazil (22° 01′S and 47°54′ W; 856 m above sea level) (Table 1). The accessions were chosen based on their origins (different ecotypes), genetic dissimilarity [60] and ploidy levels and reproduction modes. Tetraploid *P. notatum* is apomictic, whereas the diploid is sexual. The Q3664 (BGP 216) accession is a hybrid from a cross between a sexual colchicine-induced tetraploid (PT-2) and a white-stigma apomictic tetraploid of *P. notatum* [50]. The Q3664 is characterized as a facultative apomictic accession with a high level of sexuality (> 70%) [53].

For this study, five clones (biological replicates) of each accession were planted in 8 L pots, filled with 1:1 soil/vermiculite and grown in a greenhouse under the same environmental conditions. The climate is humid subtropical (according to the Köppen-Geiger classification system), with annual average low and high temperatures of 15.3 and 27.0°C, respectively, and total rainfall of 1,422.8 mm, occurring mainly during the spring and summer seasons [61].

Young leaf samples from three clones of each accession were collected during summer (December). One clone each of the accessions BGP34 (apomictic tetraploid) and BGP22 (sexual diploid) presented inflorescences, which were also collected. All leaf and floret samples were immediately placed in liquid nitrogen and subsequently stored at −80°C until RNA extraction. Total RNA was isolated according to Oliveira et al. [62]. RNA integrity was assessed in a 1% agarose denaturing gel and quantified using a NanoVue Plus spectrophotometer (GE Healthcare Life Sciences, Little Chalfont, UK).

### Verification of the mode of reproduction of *P. notatum* accessions through cytoembryological analysis

The mode of reproduction of the plants was confirmed using the embryo sac clarification method proposed by Young et al. [63], with minor modifications. Inflorescences at anthesis (when the embryo sac is fully developed) were collected and fixed in FAA (95% ethanol, distilled water, glacial acetic acid, formalin 40%, 40:13:3:3 v/v) for 24 h at room temperature. Afterward, the FAA was replaced with 70% ethanol and the samples were stored at 4°C. Ovaries were extracted by dissection under a stereoscopic microscope and stored in 70% ethanol. The embryo sacs were clarified by replacing 70% alcohol with the following series of solutions: 85% ethanol; absolute ethanol; ethanol:methyl salicylate (1:1); ethanol:methyl salicylate (1:3); 100% methyl salicylate (two times). The samples remained in each solution for 24 h. Observations were carried out with an Axiophot microscope (Carl Zeiss, Jena, Germany) using the differential interference contrast (DIC) microscopy technique. A total of 100 embryo sacs per accession were evaluated.

### Verification of the ploidy level of *P. notatum* accessions through flow cytometry

Flow cytometry analyses were performed to confirm the exact ploidy level of each plant used in the experiment and results were compared to the literature data. Approximately 5 mg of young leaf tissue from each plant was used (Table 1). *Pisum sativum* cv. Ctirad samples (2C=9.09 pg), were used as an internal control [64,65]. Samples were triturated in a petri dish containing 800 μL of LB01 buffer (0.45425 g TRIS, 0.186125 g NaEDTA, 0.0435 g-spermine, 0.29225 g NaCl, 1.491 g KCl, and 250 μL of Triton X-100 in 250 mL of distilled water, 7.5 pH, 0.11% v/v of β-mercaptoethanol) to obtain a nuclear suspension [66]. The nuclear suspension was filtered through a mesh of 40 μm and incubated at room temperature for 5 min, followed by the addition of 25 μL of propidium iodide and 25 μL of RNase. For each sample, at least 10,000 nuclei were analyzed. Samples were analyzed with a FACSCalibur flow cytometer (Becton Dickinson, New Jersey, USA). Histograms were obtained in Cell Quest software and analyzed in Flowing Software (available at http://www.flowingsoftware.com). Only histograms with both peaks (sample and standard) of approximately the same height were considered. The nuclear DNA index (pg) of the plants was estimated based on the value of the G1 peak as an internal reference. We calculated the mean value of C per biological replicate and accession and the difference between the highest and lowest values (Δ) observed in replicates of each accession.

### RNA-seq library construction, Illumina sequencing, and data quality control

The cDNA libraries were constructed from each RNA sample (18 leaf and two floret libraries) using specific barcodes and the TruSeq RNA Sample Preparation Kit (Illumina Inc., San Diego, CA, USA) following manufacturer’s instructions. Library quality was confirmed using the Agilent 2100 Bioanalyzer (Agilent Technologies, Palo Alto, CA, USA) and quantified via quantitative real-time PCR (qPCR) (Illumina protocol SY-930-10-10). Clustering was conducted using a TruSeq Paired-End Cluster Kit on a cBot (Illumina Inc., San Diego, CA, USA). Paired-end sequencing was performed on the Illumina Genome Analyzer IIx Platform with TruSeq SBS 36-Cycle kits (Illumina, San Diego, CA, USA) following manufacturer’s specifications. RNA-seq was performed with floret libraries of the accessions BGP22 (diploid/sexual) and BGP34 (tetraploid/apomictic) in two separate lanes, without biological replicates. The remaining 18 leaf libraries from six different accessions (Table 1), each with three independent biological replicates (clones), were distributed randomly in the flow cell, with three libraries per lane.

Raw data was converted to FastQ files containing 72-bp reads. Quality control was performed using the NGS QC Toolkit v2.3.3 [67]. Initially, high-quality reads (Phred quality score ≥ 20 in at least 75% of bases) and reads with more than 60 bases were selected. Subsequently, reads were trimmed at the 3’ end for the removal of barcodes. All reads were deposited in the NCBI Short Read Archive (SRA) under accession number SRP150615.

### Transcriptome assembly and completeness assessment

High-quality reads from leaves and florets were *de novo* assembled into a reference transcriptome for *P. notatum*, performed with Trinity program v2.0.2 [68] using default settings. Contigs redundancy was minimized through the selection of the first Butterfly sequence from each Chrysalis component, which is considered the most representative contig. To assess the sequenced reads-support of the assembled contigs, the high-quality short reads were mapped back to the transcriptome using the Bowtie2 sequence aligner v2.2.5 [69]. The Benchmarking Universal Single-Copy Orthologs (BUSCO) tool, an approach for the assessment of conserved orthologs among plant species in a set of sequences [70], was used to estimate the completeness of the transcriptome assembly.

### Transcriptome annotation

All transcripts were aligned to proteins from the the NCBI non-redundant (Nr) database, from the UniProtKB/Swiss-Prot and from the grass protein database from Phytozome v9.0, using the BLASTX algorithm with an e-value cutoff of 1e-06. Gene Ontology (GO) [71] terms were retrieved for transcripts showing similarity to sequences from NCBI Nr or UniProtKB databases using Blast2GO software [72]. We used REVIGO [73] to summarize and visualize GO terms sets. Pathways were assigned to metabolic pathways from the Kyoto Encyclopedia of Genes and Genomes (KEGG) database [74].

### Estimation of transcript expression levels

The expression levels of the transcripts were estimated using the FPKM method (expected number of fragments per kilobase of transcript sequence per million base pairs sequenced). Read counts for each transcript from each sample (Table 1) was obtained using Bowtie v2-2.1.0 and normalized using the RSEM software (RNA-seq by expectation maximization) [75]. A Venn diagram for the visualization of the amount of exclusive and shared transcripts among samples was created using the online platform available at http://bioinformatics.psb.ugent.be/webtools/Venn/. Principal component analysis (PCA) was used to assess expression patterns of sequenced genotypes.

### Differential expression analysis

EBSeq [76] was used to identify differentially expressed genes (DEGs) using FPKM values, at a false discovery rate (FDR) ≤ 0.05. Transcripts with a log2 fold change (FC) ≥ 1.5 in transcript abundance were regarded as overexpressed. GO enrichment analysis was carried out using R software (version 3.3.1) and the ‘goseq’ 1.24.0 Bioconductor package [77]. P-values were subjected to Bonferroni correction, and we considered adjusted p-values ≤ 0.05 as enriched. Resulting enriched GO terms were summarized using REVIGO [73].

Genotypes were grouped according to their phenotypic classes as sexual diploids (2×SEX), sexual tetraploids (4×SEX) and apomictic tetraploids (4×APO) to be compared in pairwise differential expression analyses. A two-step pipeline was used for the identification of the DEGs (Fig 6). In the first step, the objective was to select a list of equally expressed transcripts among genotypes belonging to the same phenotypic class. This procedure allowed the detection of gene expression patterns related with characteristics that were shared among genotypes, independently of physiological or developmental particularities. Thus, EBSeq was used to estimate the pairwise posterior probabilities of a transcript being equally express between genotypes: sexual diploids (accessions: BGP22 and BGP306), and apomictic tetraploids (accessions: BGP30, BGP34 and BGP115). Transcripts that presented PPEE ≥ 0.95 were kept for the second step of the analysis. The second step consisted of another round of pairwise differential expression analyses between phenotypic classes: (i) 2×SEX vs. 4×APO; (ii) 2×SEX vs. 4×SEX; (iii) 4×APO vs. 4×SEX.

**Fig 6.**
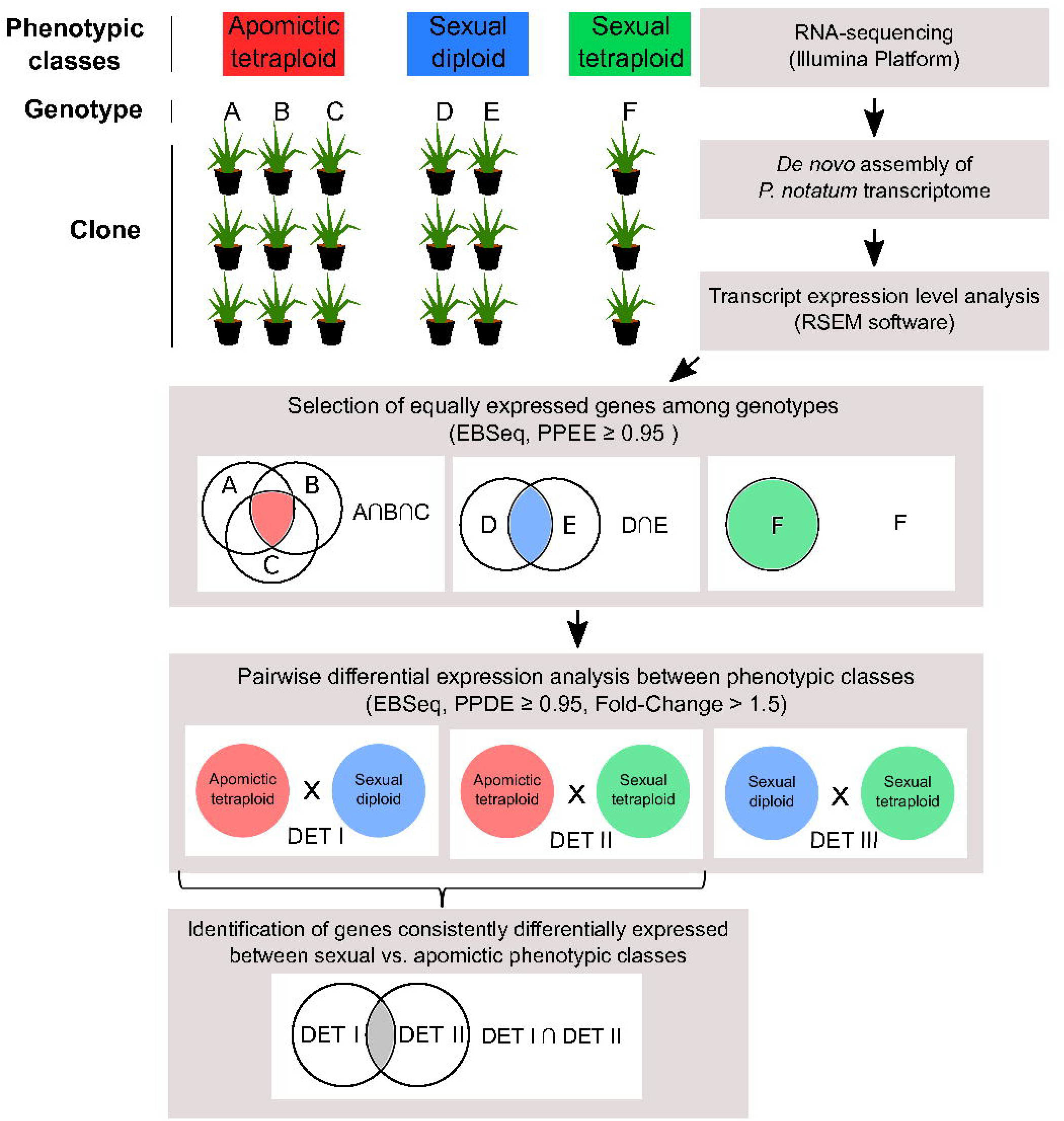
Workflow of differential expression analyses used in this study. We realized pairwise differential expression analyses of all three phenotypic classes (apomictic tetraploid “4×APO”, sexual diploid “2×SEX” and sexual tetraploid “4×SEX”). Each phenotypic class had different genotype with three clones. In the first step, the objective was to select a list of equally expressed transcripts among genotypes belonging to the same phenotypic class. EBSeq was used to estimate the pairwise posterior probabilities of a transcript being equally express (PPEE) between genotypes of the same phenotypic class: apomictic tetraploid (A×B; A×C; B×C) and sexual diploid (D×E). Transcripts that presented PPEE ≥ 0.95 were kept for the second step of the analysis. The second step consisted of another round of pairwise differential expression analyses between phenotypic classes: (i) 4×APO vs. 2×SEX; (ii) 4×APO vs. 4×SEX; (iii) 2×SEX vs. 4×SEX. The objective was to select a list of transcripts that presented pairwise posterior probabilities of being differentially expressed (PPDE ≥ 0.95). Additionally, we identified a list of transcripts that were consistently differentially expressed between sexual and apomictic samples, independently of the ploidy level (diploid or tetraploid).

### Quantitative reverse transcription PCR (qRT-PCR) validation of differential expression results

To verify the reliability and accuracy of transcriptome data and differential expression analyses pipeline, 18 DEGs were randomly selected for quantification through reverse transcription PCR (qRT-PCR). Eight transcripts showing similar expression patterns based on FPKM values were used as internal reaction controls. Primer sets for controls and targets were designed using Primer3Plus software [78]. All primer pairs were initially tested in regular PCR reactions using genomic DNA as a template from the same genotypes and clones used in RNA-seq. Only primers that amplified the genomic DNA of all genotypes were used in the following qRT-PCR amplification efficiency tests. Primer pairs with an amplification efficiency of 90-110% and R^2^> 0.99 were used for relative expression analyses.

Total RNA (500 ng) was used for cDNA synthesis using the QuantiTect Reverse Transcription Kit (Qiagen Inc., Chatsworth, CA, USA). qRT-PCR were performed using a CFX384 Real-Time PCR Detection System with iTaq Universal SYBR Green Supermix (Bio-Rad Laboratories Inc., Hercules, CA, USA), according to the manufacturer’s instructions. The reaction conditions were 95°C (10 min) and then 40 cycles of 95°C (30 s) and 60°C (1 min). All experiments were performed using technical triplicate, and no-template controls were included. The specificity of amplicons was confirmed through the analysis of the melting curve. The baseline and Cq (quantification cycle) values were automatically determined, and the expression values were estimated using the ΔΔCt method implemented by the CFX Manager 2.1 software (Bio-Rad Laboratories, Inc., USA). Reference genes were selected according to their expression stability (M < 0.5) and coefficient of variance (CV < 0.25) among samples. Mann-Whitney U-tests were performed to estimate statistical significance of distribution differences between samples.

### Search for putative apomixis genes

To perform a detailed comparative analysis of *P. notatum* transcripts obtained in this study with their syntenic counterparts on rice chromosomes that are conservatively linked to apomixis in *Paspalum* species [8], we aligned all *P. notatum* transcripts to rice transcripts using the BLASTN tool and selected those that presented putative homology to genes in the highlighted area of rice chromosomes 2 and 12. Then, we selected only *P. notatum* transcripts present in the ACR, which were delimited by a set of molecular markers completely linked to the apospory locus in this species. Thus, the region encompassed the C1069 and C996 markers for rice chromosome 12 and between C560 and C932 markers for chromosome 2 [8,21,24-26,79]. We also compared the *P. notatum* reference transcriptome with sequences reported for the genus *Paspalum*, potentially associated with apomixis. The sequences were used as queries for BLASTN search in the 114,306 transcripts database, with an e-value cutoff of 1e-05.

### Construction of a transcripts coexpression networks

To generate coexpression network, we used expression values in FPKM of all assembled transcripts from floret and leaf tissues. Transcripts showing null values for most of the replicates were excluded to reduce noise and eliminate residuals in the analysis. We calculated an all-versus-all coexpression network matrix using the Pearson correlation coefficient cutoff of ≥ 0.8. The highest reciprocal rank (HRR) method proposed by Mutwil et al. [80] was used to select only the strongest edges, considering an HRR limit of 30. The Heuristic Cluster Chiseling Algorithm (HCCA) was applied to partition the networks into manageable clusters, with three-step node vicinity networks (NVN) [80]. The interactive coexpression network was visualized using the Cytoscape 3.4.0 software [81].

## Acknowledgments

We are grateful to Maria Augusta C. Horta and Natalia F. Murad for their help in generating the coexpression networks and Mariana V. Cruz for critically reading the manuscript.

## Supporting Information

**S1 Fig. Length distribution of the number of non-redundant transcripts successful annotated.** Comparison of the all non-redundant transcripts and the number of transcripts annotated in the NCBI non-redundant protein database by size range.

**S2 Fig. Principal component analysis (PCA) according to the FPKM values of *P. notatum* transcriptome.** PCA analysis of leaf and floret transcriptomes for all genotypes and clones used in RNA-seq.

**S3 Fig. Histograms of gene expression obtained by qRT-PCR.** qRT-PCR validation showing the relative expression patterns of 4 genes that are differentially expressed between tetraploid apomictic "4×APO" (red box) and diploid sexual "2×SEX" (blue box). *P < 0.05; **P < 0.01; ***P < 0.001: statistically significant differences in gene expression between phenotypic classes compared using the Mann-Whitney U-test.

**S4 Fig. Histograms of gene expression obtained by qRT-PCR.** qRT-PCR validation showing the relative expression patterns of 4 genes that are differentially expressed between tetraploid apomictic "4×APO" (red box) and tetraploid sexual "4×SEX" (green box). *P < 0.05; **P < 0.01; ***P < 0.001: statistically significant differences in gene expression between the phenotypic classes compared using the Mann-Whitney U-test.

**S5 Fig. Functional classification of enriched overexpressed DEGs in the 2×SEX vs. the** 4×**APO comparison**. Gene Ontology biological process (blue boxes), Gene Ontology cellular component (yellow boxes), and Gene Ontology molecular function (orange boxes). a) Categories enriched in overexpressed transcripts in 2×SEX and b) categories enriched in overexpressed transcripts in 4×APO.

**S6 Fig. Functional classification of enriched overexpressed DEGs in the 2×SEX vs. 4×SEX comparison.** Gene Ontology biological process (blue boxes), Gene Ontology cellular component (yellow boxes), and Gene Ontology molecular function (orange boxes). a) Categories enriched in overexpressed transcripts in 2×SEX and b) categories enriched in overexpressed transcripts in 4×SEX.

**S7 Fig. Functional classification of enriched overexpressed DEGs in the 4×APO vs. 4×SEX comparison.** Gene Ontology biological process (blue boxes), Gene Ontology cellular component (yellow boxes), and Gene Ontology molecular function (orange boxes). a) categories enriched in overexpressed transcripts in 4×APO and b) categories enriched in overexpressed transcripts in 4×SEX.

S1 Table. Primer sequences and amplicons of the candidate reference genes evaluated in this study.

S2 Table. Quantitative RT-PCR primer sequences.

S3 Table. BLAST search results for sequences of *P. notatum* from the literature against the assembled transcriptome.

